# Sustainable use of plant protection products in heroic viticulture areas. Reducing risks to the environment and winemaking

**DOI:** 10.1101/2021.07.16.452637

**Authors:** J. Antonio Cortiñas, M. Eva Fernández-Conde

## Abstract

**Objective:** The aim and objective of this article is to focus on the importance of the rational use of plant protection products to combat vine fungi and at the same time reduce their risks to human health and the environment. The integrated management of fungal diseases aims to achieve the development of healthy crops with the minimum alteration of agro-ecosystems and the promotion of natural mechanisms.

**Material and methods:** The study was carried out in one vineyard in Ribeira Sacra (North-West Spain) during 2018. Souto vineyard is located at 438 m above mean sea level (42° 24’ 27.67” N 7° 28’ 20.06” W; northwest-southeast orientation) in the lower terraces of the river Sil’s banks, following the contour lines and with gradients of up to 80%. The variety studied was Godello, for sampling the reproductive structures in the air (spores of *Botrytis* and *Erysiphe* and sporangia of *Plasmopara*), a Lanzoni VPPS-2000^®^ spore trap (Lanzoni s.r.l., Bologna, Italy) was used.

**Results:** The *Botrytis* Seasonal Spore Integral (SSIn) was markedly higher than for the other pathogens under consideration. Taking into account the maximum daily values, a clear dominance of *Botrytis* spores was also found, with a maximum of 397 spores/m^3^ at the beginning of June, while *Erysiphe* and *Plasmopara* were recorded at around 26 and 227 spores/m^3^, respectively, at the beginning of august and mid-July. The statistical analysis of the spore concentrations and the main meteorological variables showed for *Erysiphe* that the highest Spearman’s **r** correlation coefficient corresponded to the rainfall, as for *Plasmopara* airbone sporangia, but with a negative sign, while for *Botrytis* spores, no significant correspondence was found for any meteorological parameter.

**Conclusion:** The use of plant protection products can be much more effective if fungicides are applied at the right time, at the precise doses and combined with agricultural techniques of management of the vineyards. There are sustainable and profitable alternatives that can improve vine yields while protecting the environment in areas of heroic viticulture where the vineyard, is a fundamental element of the wine-growing landscape, combining as it does historical, cultural and landscape characteristics.

## 1. Introduction

The vineyard in the Ribeira Sacra dates backs to the Roman times. The initial expansion of the cultivation is linked to the presence of numerous monasteries in the area (Hidalgo, 2011). Nonetheless, the viticulture of the zone initiates the current development from the creation of the Regulatory Council of the Ribeira Sacra Designation of Origin at the end of the S. XX (C.R.D.O.R.S., 2017). The regulation of the C.R.D.O.R.S. fixed the territorial limits, the cultivable varieties and all organoleptic characteristics which typify this wine.

Few studies have addressed the wine-making potential of this area, or the varieties currently grown there, although there has been some research into the genetic characterisation of certain grapevine cultivars and into the development of systems to confirm the authenticity of local protected designation of origin wines (Rebolo et al. 2007).

The Ribeira Sacra landscape, on which this study focuses, is particularly representative of a model of grape production that depends on specific factors linked to the physical environment, and the socio-economic orientation of the area. Although viticulture practices can reduce the harmful effects of pathogenic fungi they cannot be completely avoided (Guerin-Dubrana et al. 2019).

In this area where the vineyard is a fundamental element of the landscape and where all the field work is done by hand, it is important to make a rational use of phytosanitary products in order to increase the profitability of the winegrower by reducing costs in their application and at the same time reducing their impact on the environment.

Traditionally, cryptogamic diseases have been combated by means of chemical products, using dates on a calendar, generally involving increased doses in times of heat and high humidity. However, these treatments tend to be excessive and are often not applied at the right time.

In this context, the study of phytopathogenic fungi affecting vineyards is not only of commercial interest but also a means of protecting the natural heritage that these areas represent.

Food safety and security is an issue of growing interest. Consumers are exposed daily to potentially toxic substances in food such as residues of plant protection products from various practices used in the production, preparation, conservation and handling of food (Camean and Repetto, 2006).

The objectives of the present study are therefore:

- To analyse the incidence of the main cryptogamic diseases affecting the Godello variety in the Souto plot.
- To review the type of antifungal treatments used.
- To offer sustainable and profitable alternatives that improves grape harvests while protecting the environment.

## 2. Material and methods

The study was carried out in one vineyard in Ribeira Sacra (NW Spain) (Figure 1). To ascertain the climatic characteristics, the meteorological data were obtained from the Xábrega meteorology station (www.meteogalicia.gal). The year 2018 was very wet (814.4 mm), with March standing out with a total of 310.1 mm collected, while the driest month was August with only 3.0 mm.

**Figure 1.**
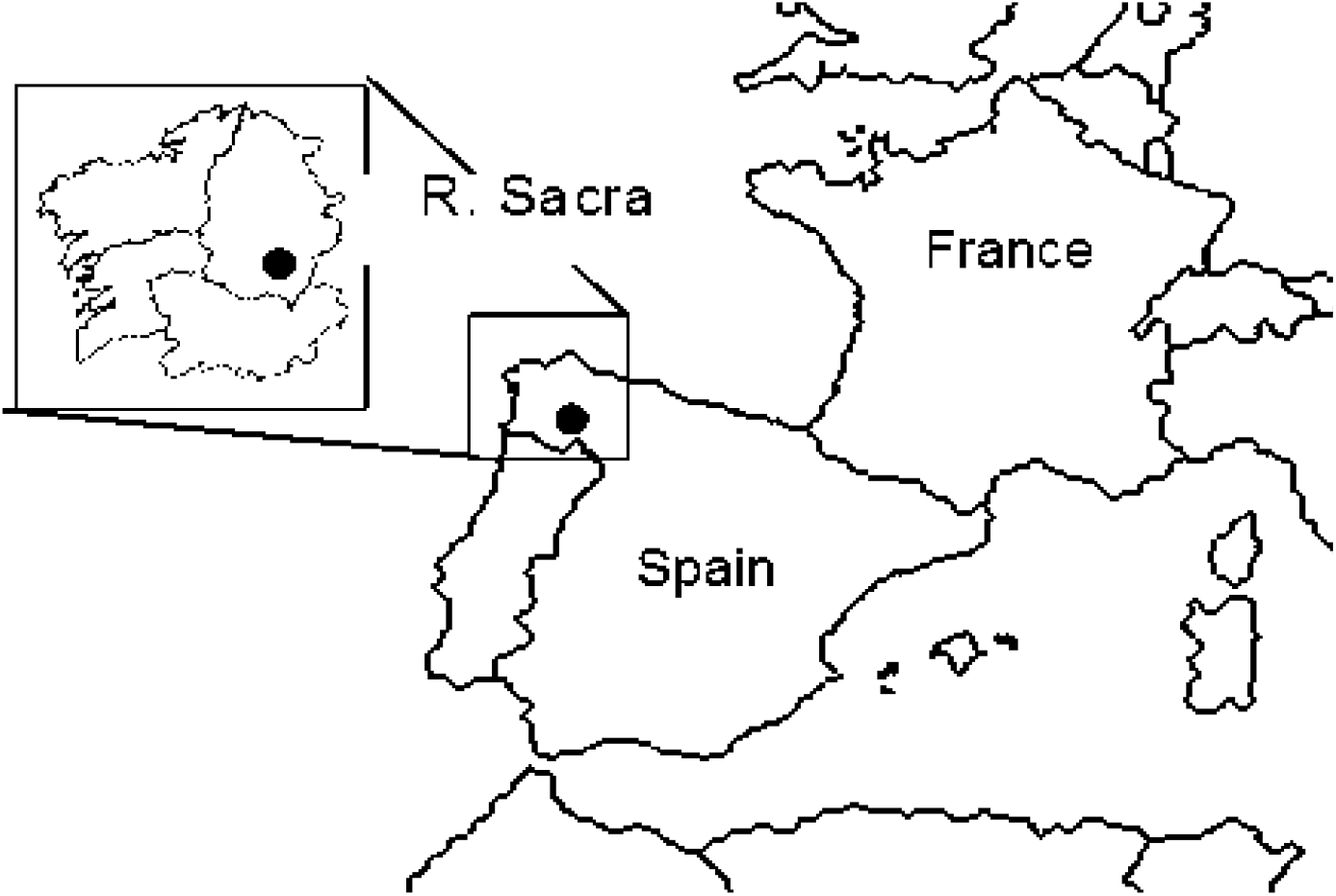
Location of Souto plot in the Ribeira Sacra region of northwest Spain

The average maximum and minimum temperature was (31.5 °C max T^<u>a</u>^ and 14.5 °C min T^<u>a</u>^). Average relative humidity was of 77.1%. Thus, in the warmest month, temperatures in the terraces closest to the river can be as much as one and a half degrees higher than the average for Ribeira Sacra as a whole.

The phytopathological study focused on the most widespread diseases in the vineyards of the area, grey rot caused by *Botrytis cinerea*, powdery mildew caused by *Erysiphe necator* and downy mildew caused by *Plasmopara viticola*. For sampling the reproductive structures in the air of the three fungi, a Lanzoni VPPS-2000^®^ spore trap (Lanzoni s.r.l., Bologna, Italy) was used (Hirst, 1952) (Figure 2).

**Figure 2.**
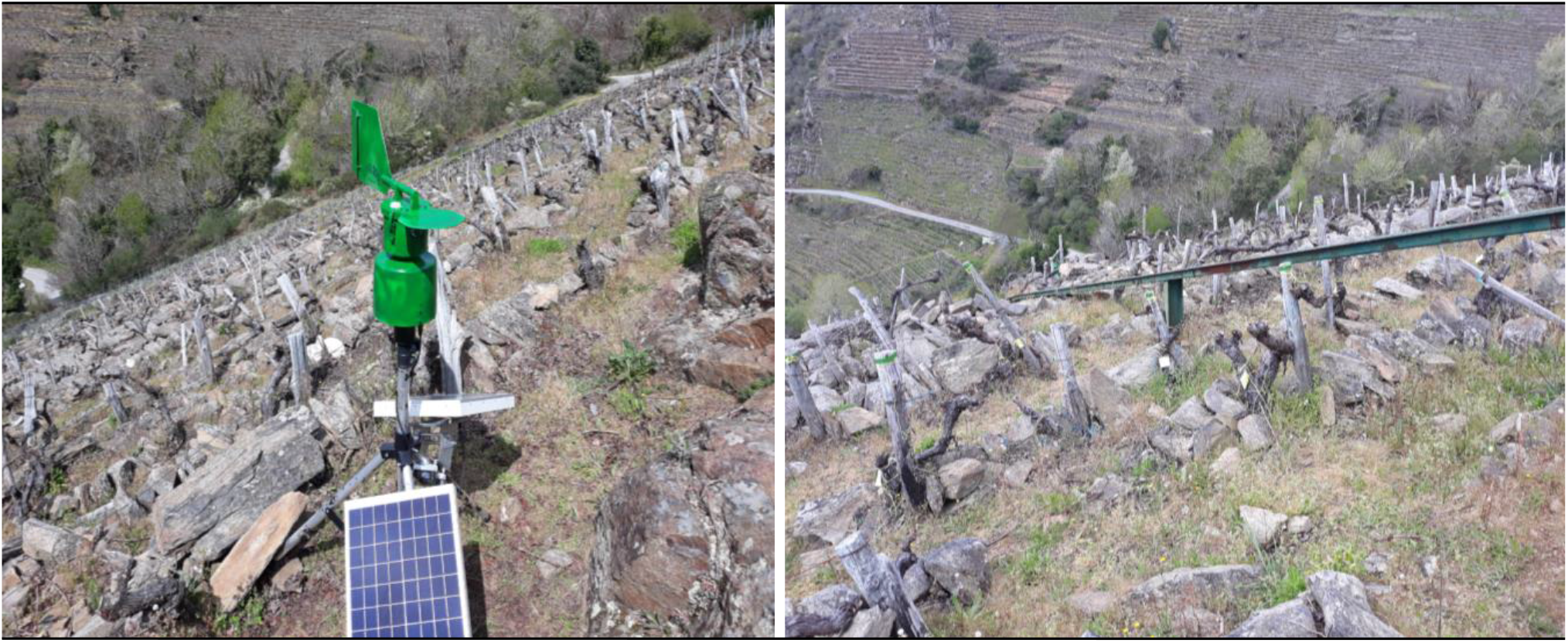
Lanzoni VPPS^®^ spore trap in the Souto plot.

The volumetric sampler was placed at a height of two metres, according to the leaf layout of the vine. For the identification and count of spores and the preparation of samples the protocol proposed by the Spanish Network of Aerobiology was followed (Galán et al. 2007). The counts were made in two longitudinal transects along the slides and the concentrations were expressed as a daily average of fungal spores/m^3^ of air. This study was carried out over the course of the entire active grapevine cycle.

To complete the phytopathological study, a phenological control was also carried out using the scale proposed by the BBCH (*Biologische Bundesanstalt für Land - und Forstwirtschaft, Bundessortenamt und CHemische Industrie*). The cultivation system in Souto forms a tall goblet, with three arms.

## 3. Results

The *Botrytis* Seasonal Spore Integral (SSIn) was markedly higher than for the other pathogens under consideration, with a maximum of 4,910 spores, while *Erysiphe* and *Plasmopara* recorded 468 and 1,354 spores, respectively (Table 1; Figure 3).

**Table 1.**
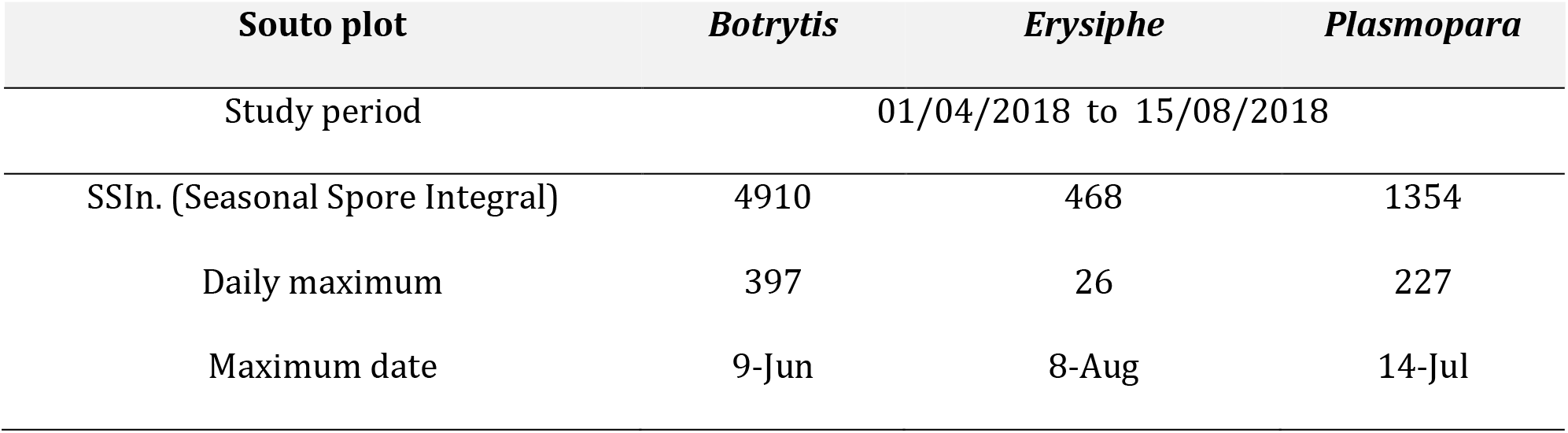
SSIn data, daily maximum spore concentration and maximum date for *Botrytis*, *Erysiphe* and *Plasmopara* in Ribeira Sacra (Souto plot) during the study period (spores/m^3^ of air).

**Figure 3.**
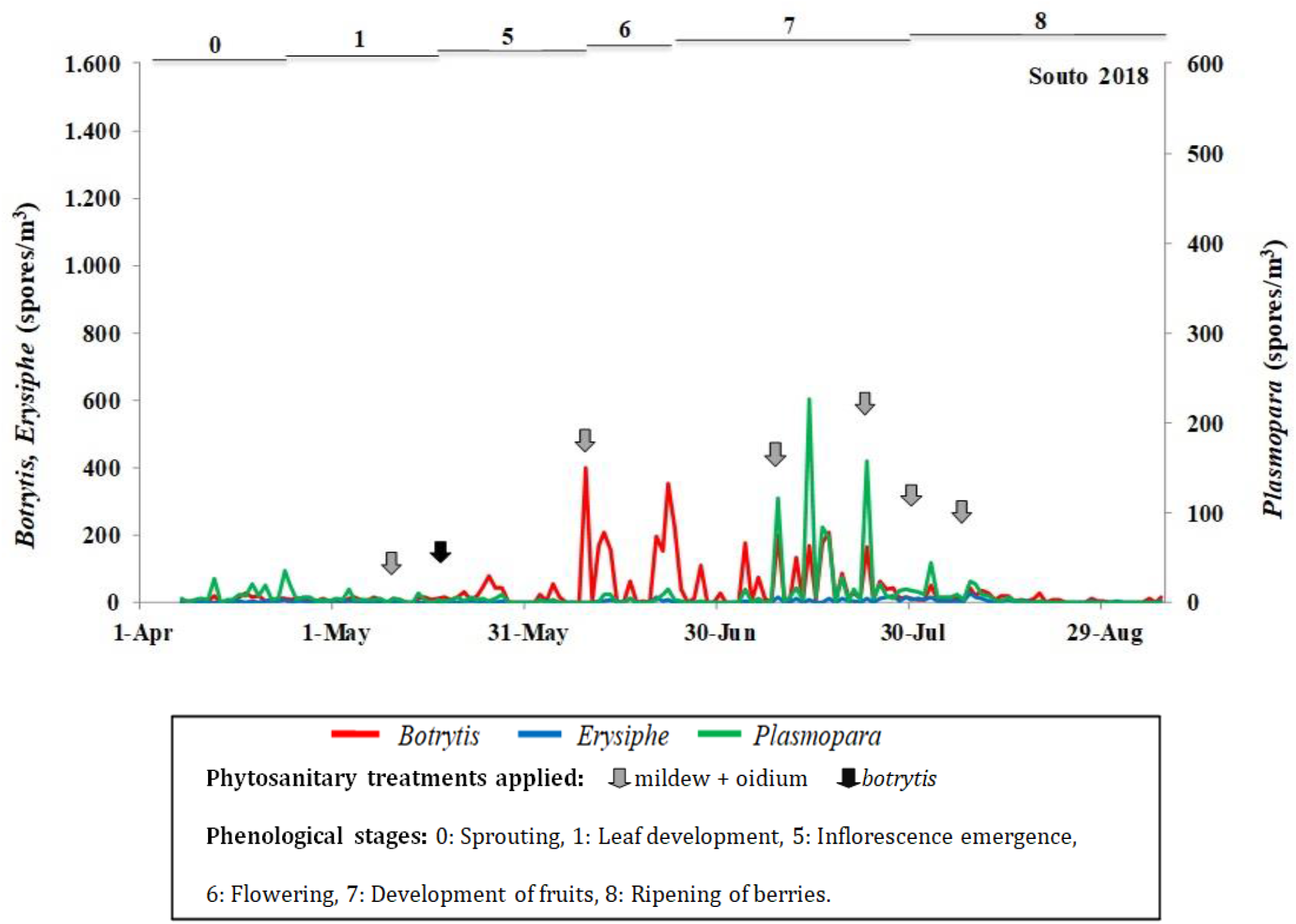
Phenology, spore concentrations, and phytosanitary treatments applied in the Souto plot (Godello variety) Ribeira Sacra.

Taking into account the maximum daily values, a clear dominance of *Botrytis* spores was also found, with a maximum of 397 spores/m^3^ at the beginning of June, while *Erysiphe* and *Plasmopara* recorded 26 and 227 spores/m^3^, respectively, in August an July. The *Botrytis* and *Erysiphe* airborne spore presence remained almost constant throughout the grapevine growth cycle, whereas *Plasmopara* did not exhibit a continuous daily record. In terms of the pathogens’ relationship with the various phenological phases of the grapevine, the highest *Botrytis* incidence was detected from the inflorescence emergence (Stage 5) stage until the end of flowering (Stage 6). To complete the phytopathological study, a phenological control was also carried out using the scale proposed by the BBCH (*Biologische Bundesanstalt für Land - und Forstwirtschaft, Bundessortenamt und CHemische Industrie*), from the sprouting of the vines to the harvesting of the grapes.

A statistical analysis comparing the daily spore concentrations and the daily values of the main meteorological variables throughout the grapevine reproductive cycle was conducted, also taking into account the daily values of the meteorological variables during the previous seven days in terms of the presence of spores in the atmosphere of the vineyard. The statistical analysis of the spore concentrations and the main meteorological variables showed that rainfall and relative humidity had a statistically significant influence, but negative. Within the significant *Erysiphe* correlations, the highest Spearman’s r correlation coefficient was found for rainfall (−0.274**) and for *Plasmopara* airborne sporangia was also rainfall (−0.270**). For *Botrytis* spores, no significant correspondence was found for any meteorological parameter (Table 2).

**Table 2.**
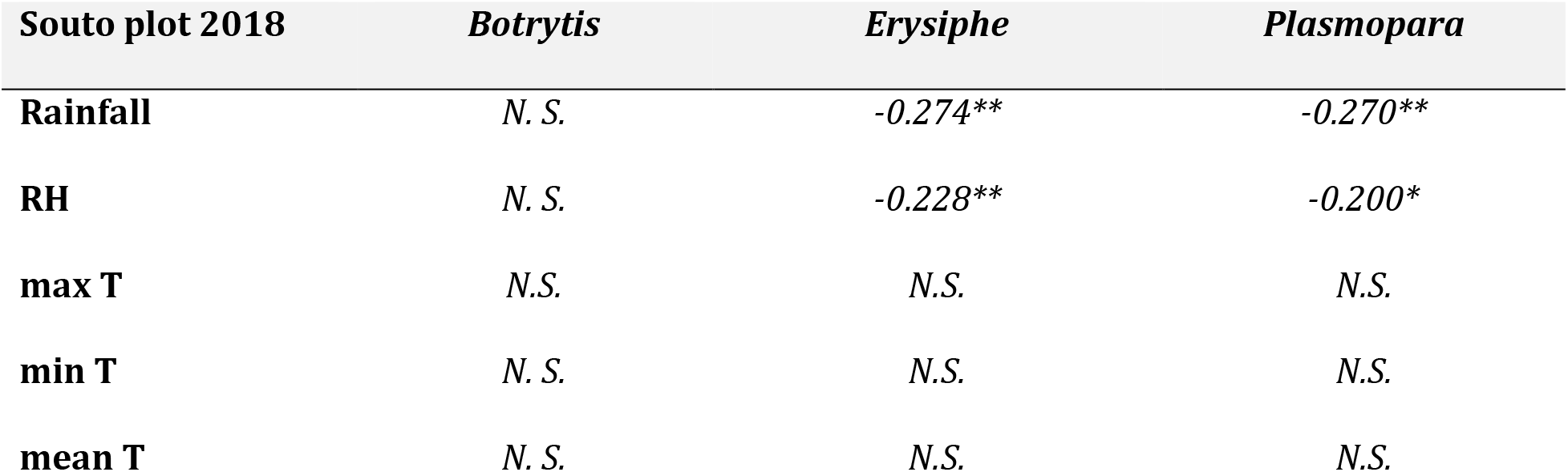
Spearman’s rank correlation of different parameters for the studied cultivars. Spore concentrations (*Botrytis*, *Erysiphe* and *Plasmopara*) and the main meteorological parameters. (*plevel*: * < 0.05; ** < 0.01; N.S.: not significative).

| Souto plot 2018 | *Botrytis* | *Erysiphe* | *Plasmopara* |
| --- | --- | --- | --- |
| **Rainfall** | *N. S.* | -0.274** | -0.270** |
| **RH** | *N. S.* | -0.228** | -0.200* |
| **max T** | *N.S.* | *N.S.* | *N.S.* |
| **min T** | *N. S.* | *N.S.* | *N.S.* |
| **mean T** | *N. S.* | *N.S.* | *N.S.* |

## 4. Discussion

The chemical fungicide application following preset treatment schedules is a widely used control strategy among wine growers to reduce the fungal disease impact on the crop, which has several environmental repercussions (Gargallo et al. 2018). Aerobiological models associated with climatic variables and the use of local cultivars, adapted to the specific area conditions, would enhance the control of grey mould, powdery mildew, and downy mildew (Casanova, 2003).

Grey mould (*Botrytis*) affects over 200 mainly dicotyledonous species and has a serious economic impact, especially on the grapevine (Kassemeyer and Berkelmann-Löhnertz, 2009). In addition to the loss of crop yields in harvests, it can also reduce the quality of the wine, leading to an unstable colour, unpleasant tastes and difficulties with clarification (Aleixandre et al. 2013).

*Botrytis* was the pathogen with the highest concentrations recorded in this study. Although *Botrytis* infections have been detected in all phenological stages, flowering appears to be more conducive to its development (Molitor et al. 2016). The influence of the pathogens has been corroborated with the analysis of the collected samples in the field.

Temperature and rainfall are two determining factors in the development of *Botrytis* infection (Oliveira et al. 2009), with the optimal temperatures for infection being around 18-20° C (Latorre et al. 2002). For its part, rain may influence airborne spore concentration both positively, by triggering spore release, and negatively, by removing fungal spores from plants by means of rain-out and wash-out effects (Pace et al. 2019). Both factors would explain the higher concentration of *Botrytis* in the summer season, coinciding with flowering and the beginning of fruit setting (May-July).

Some authors noted that the critical period for *Erysiphe* infection ranges from the beginning of flowering until the berries have a diameter of 7 mm or reach 8 °Brix (Campbell et al. 2007). Temperature is one of the factors most influencing the development of this fungus, the optimum temperature being estimated at around 25°C; infection is reduced at temperatures of below 8°C and above 33°C (Calonnec et al. 2004). In this study, the concentrations of the fungus were very low during the whole vegetative cycle of the vine, without any remarkable peak.

At the beginning of August, the concentration of spores in the air is still much lower, related to the increase of temperatures during this month until the first days of September. Along the studied period, the maximum temperature registered a value higher than 33 °C in more than 30 days.

Grapevine downy mildew (*Plasmopara*) is native to North America and was first detected in Spain in 1880. Of the fungal spores counted in the present study, *Plasmopara* exhibited average values occupying an intermediate point between *Botrytis* and *Erysiphe* (Table 1). In the distribution pattern of atmospheric spores, an early peak in April coincided with leaf development (S1), although its maximum concentrations are usually located from the development of fruit onward. Previous studies carried out on red varieties grown in Ribeira Sacra offer similar data to those obtained in the present study in terms of the incidence of the disease as estimated by the concentration of total spores (Cortiñas et al. 2020).The temperature determines the length of the ripening period of this fungus, the optimum temperature range being between 20 and 25 °C (Barrios and Reyes, 2004). In the plot studied, when the disease became a major problem a significant increase in the maximum temperature and rainfall over a certain period of time was observed, something that would tend to favour the germination of spores and the spread of the disease. Humidity is another important climatic parameter in secondary infections (Pérez-Marín, 2004). Optimal values range from 95 - 100% and problems for reproduction if humidity falls below 75%. Indeed, with low values of relative humidity and high temperatures, *Plasmopara* spores find it difficult to survive (Gessler et al. 2011). During spring, the humidity was relatively low, rain and fog was sporadic and average temperatures generally remained below 15 °C, coinciding with a low incidence of downy mildew.

### 4.1. Assessment of phytosanitary treatments applied in the vineyard

The phytosanitary treatments applied have been decided by the winegrowers, following the same guidelines as usual in this geographical area. Therefore, the study of pathogens in the crop was carried out independently of the treatments applied. Thus, at the end of the harvest and by comparing the results of the experimental study, it was possible to estimate its effectiveness both by the type of phytosanitary treatment and by the time of application. Two different types of phytosanitary treatments were applied: against *Botrytis* (one application) and one combined against mildew and oidium (six applications). In all cases, the fungicides used until flowering (S6) were systemic, and from this moment onward of the contact type (mainly sulphur-based powder products until the fruit set, and copper in wet dust until the softening of berries). The treatment against *botrytis* was applied in leaf development (S1) and considering that spore concentrations of this pathogen in air were very low, this did not seem necessary. However, in phenological stages of inflorescence emergence (S5) and flowering (S6) there were peaks of this fungus and no treatment was applied.

The combined products against both mildew and oidium by contrast were administered much more frequently, with a total of 6 treatments. The treatments applied during development of fruit (S7) were necessary, in the present authors’ opinion; however, the subsequent treatment during the ripening of berries was harder to justify, due to the low incidence of both pathogens at this stage. The treatments applied, all of them had a clear impact in terms of reducing the fungal spores/m3 in the air, so the authors regard them as useful, except the treatment administered during the leaf development stage (S1), where the low incidence of the pathogens did not justify the application.

No specific treatments against mildew were administered in Souto plot, the vineyard being protected at the moments of greatest incidence of the fungus by the combined treatments applied against both mildew and oidium.

### 4.2. Alternatives for a more sustainable control of phytosanitary products in Ribeira Sacra

In general it is advisable to use chemical controls only as a last resort, while systemic fungicides should be discarded with whenever possible because of their likelihood of creating resistance within the fungus (Cortiñas et al. 2020). Contact fungicides such as sulphur or copper are preferable, even though they only protect preventively; these are low-cost products with a preventive effect whose use in vineyards has been known for more than 100 years (Alti et al. 2016).

If crop and weather conditions determine the use of systemic fungicides, we must use different active principles, in order to reduce the probability of generating resistance in the fungus. The correct doses of application and the right moment will be determined by the epidemiological study of the pathogen.

Climate information provided by local weather stations, as well as aerobiological information models together with native varieties better adapted to each geographical area, will allow a better and more exhaustive control of vineyard diseases (Pérez-Sanz et al. 2008).

Phytosanitary alert systems based on weather conditions that are conducive to the development of spores allow for a much more reasonable adjustment of the application of phytosanitary treatments.

Other alternatives to the traditional use of chemical products are those related to biological control. In this sense we have, among other possibilities, the selected strain of the bacterium *Bacillus subtilis* and its capacity to produce lipopeptide antibiotics that prevent the pathogenic spores of the plants from germinating, altering the growth of the germ tube of the spores and thus inhibiting the fixation of the plant pathogen to the leaf. This biofungicide is highly environmentally friendly and broad-spectrum with extraordinary biocontrol activity against powdery mildew, downy mildew and *botrytis*, among other fungi that attack the vineyard. Lacherre and Ruíz (2014) also reported the antagonistic effect of *Clonostachys rosea* on *Botrytis*, which could also be reduced by *Aureobasidium* and *Ulocladium* (Köhl et al. 1999).

The alternative to the exclusive use of synthetic chemical plant protection in the fight against powdery mildew should be focused on three areas: preventive agronomic treatments, biological control and chemical fungicides used rationally (Cortiñas et al. 2020).

An alternative to biological control of powdery mildew is AQ-10 (Ecogen, Inc, USA). This is a fungicide that stands out for its mode of action (hyperparasitism), affecting all stages of development of the pathogen. Furthermore, it does not present any phytotoxicity risk nor does it affect the must fermentation so that the natural aromas of the wine grapes are not altered, it does not have a safety period for the cultivation and it does not generate residues in the fruit.

Based on knowledge of *Plasmopara* behaviour and in accordance with the meteorological conditions of the environment, different monitoring models have been developed covering different phases of its biological cycle. The first takes into account the spore ripening period in winter, where a qualitative and quantitative assessment is made of the degree of the spores’ maturity and the aggressiveness of the first contamination. This produces the Potential State of Infection or PSI model (Rocafort, 2015). To monitoring the length of the incubation period, the Goidanich model is used, followed for many years by the Agricultural Alert Stations with the aim of introducing Targeted Fighting, which has made it possible to rationalise the implementation of treatments by being able to focus them at the right times and reduce their number to the essential minimum.

In order to follow the evolution of the fungus, once the first infection from the wintering oospores has been observed, it is necessary to know the average temperature (mean T °C), the average relative humidity (mean RH %) and the rainfall on a daily basis. Goidanich presents a table of daily evolution in which, for each temperature, a daily growth of the fungus is fixed according to the (mean RH %) is high or low. The author defines that the (mean RH %) is high on cloudy days with low thermal difference, and at the same time expresses as low (mean RH %) the one produced on calm days with high thermal difference. Thus, the table is used in which for each (mean T °C) there are two columns, one when the (mean RH %) is lower than 75% and the other when it is higher than 75%, providing, therefore, two numerical values of daily development for the same temperature according to the (mean RH %).

When contamination occurs due to rain close to or greater than 10 mm, from the following day onwards the daily growth assessment begins, which will be added up every day until the value of 100 is reached, at which point the theoretical incubation period ends and the contamination of the fungus is evident by the appearance of the oil spots and asexual fruiting.

Another important aspect is to reduce the use of herbicides to control adventitious vegetation. In Ribeira Sacra the thatched roofs are one of the most used solutions to control the proliferation of adventitious vegetation without resorting to herbicides. Improved access to the vineyard areas facilitates the transport of the bulky straw packs to the vineyards, where they are then spread. This is a laborious and necessarily manual job, but this artificial plant cover can act as a natural herbicide for a minimum period of two years.

The accumulation of herbicide residues and other phytosanitary treatments can become a particularly serious environmental problem in areas of heroic viticulture such as Ribeira Sacra. Due to the shallowness of the soil, the residues tend to reach high concentrations over time, and washing out by rain is much less than in other types of soil. The substances that kill off the adventitious vegetation also eliminate microorganisms that are beneficial to the crop. Unwanted vegetation was traditionally fought with the tilling of the vineyard, which is totally unfeasible nowadays due to the lack of labour and the unbearable costs it would entail. Some wineries also use small motorised cultivators equipped with milling machines to work the land in some of their plots, but this does not seem to be a solution applicable to vineyards set up on narrower terraces, thus significantly increasing costs. One of the objectives is therefore to maintain permanent plant cover on those plots where any type of mechanisation is unfeasible. The disadvantage of this formula is that the vegetation must be controlled periodically so that it does not proliferate excessively and compete with the vine, which means a high need for labour by the wineries and a significant increase in production costs.

In recent years, different solutions have been assessed to eradicate or at least reduce the use of herbicides in the vineyards of the Ribeira Sacra. One of the most curious was the use of the remains of the industrial cut of the slate for the soil coverings. However, the experts did not follow this line of research because they understood that this material could significantly alter the soil conditions and the composition of the wine. Tree bark is another option proposed to control unwanted vegetation. It should be borne in mind that all the alternative systems to herbicides have their pros and cons, and there is no formula that is completely valid in all respects.

Lastly it should be noted that the vast majority of vine varieties grown in the world are highly sensitive to fungal diseases, which is why work is also being done to obtain hybrid varieties, obtained from crossing traditional and American grapevines that exhibit resistance to these diseases (Ruíz-Garcia, 2015).

## Conclusions

There are sustainable and profitable alternatives that can improve vine yields while protecting the environment in areas of heroic viticulture where the vineyard, in addition to its economic importance as a crop, is a fundamental element of the wine-growing landscape.

Depending on the epidemiological development and the concentration of the inoculum of the pathogenic fungi, it is possible to determine the type of phytosanitary product required in each case and ascertain the frequency of its application. The study of the airborne spores concentrations in the vineyard is of great interest to establish the correct guidelines for preventive treatment.

As a final conclusion, the sustainable wine-growing model would appear to be perfectly viable in the Ribeira Sacra D.O. area. In terms of methodology, the multidisciplinary research reported here served to strengthen the conclusions obtained.

## Acknowledgments

The authors thank to the Xunta de Galicia (Consellería de Educación, Universidade e F. Profesional), and Economy and Competence Ministry of Spain Government.

